# Engineering magnetically guided bacteriophages for precision antimicrobial therapy and targeted biofilm eradication

**DOI:** 10.1101/2025.10.05.680512

**Authors:** Yeping Ma, Sylvia Yang Liu, Minqi Hu, Wei Wei, Kaman Ma, Borui Zhu, Sirui Li, Bee Luan Khoo, Song Lin Chua

## Abstract

Viruses, including bacteriophages, rely on passive diffusion to reach their hosts, limiting the efficacy of virus-based therapies and leading to off-target accumulation with systemic effects. Here, we present a strategy to confer controllable motility to bacteriophages by incorporating iron nanoparticles (FeNPs) into their head structures while preserving infectivity. Cryo-electron microscopy (Cryo-EM) and transmission electron microscopy-energy dispersive spectroscopy (TEM-EDS) confirmed the FeNP presence in the phage’s head. FeNP-tagged phages can be magnetically enriched and isolated from bacterial prey cultures, eliminating the need for ultracentrifugation. Under magnetic guidance, these engineered phages exhibit rapid and directed movement through complex microenvironments, including mazes and polymer barriers, enabling precise bacterial targeting and biofilm eradication in a microfluidic system. In an *in vivo* wound infection model, magnetically guided phages successfully navigated from the peritoneum to the biofilms on the wound, thus selectively eliminating biofilms while minimizing systemic exposure to other organs. Hence, our approach confers viral mobility, thus enhancing the precision and efficacy of bacteriophage-based biotechnological and therapeutic applications.

## Introduction

Viruses, including bacteriophages, lack active motility and rely entirely on passive diffusion to encounter their specific hosts (White et al., 2021). While recent studies showed that influenza virus exhibited limited short-range motions across the host cell surface by binding onto host cell receptors with its hemagglutinin and neuraminidase (Sakai et al., 2018), viruses typically rapidly disseminate from an infection site to distant parts of the host body by ‘hitching’ a ride on the host circulatory system (Gaudin and Goetz, 2021), or spread via cell-cell contact transmission (Zhong et al., 2013). This fundamental limitation constrains the efficiency of virus-based therapeutics, particularly in phage therapy, oncolytic virotherapy, and viral gene delivery, where precise targeting to target tissues or penetration into tissues is essential (Zheng et al., 2019). The inability to control viral transport often results in off-target accumulation, systemic dispersal, and diminished therapeutic efficacy (Butt et al., 2022; Singh et al., 2024). This challenge is especially pronounced in bacteriophage therapy, where infections often occur within structured environments such as biofilms, which serve as protective niches that limit phage penetration and bacterial eradication (Bond et al., 2021; Simmons Emilia et al., 2020). Current strategies to enhance phage delivery, such as chemical modifications or encapsulation (Loh et al., 2021), often compromise infectivity, stability, or host specificity, underscoring a critical need for new approaches to actively guide viruses toward their intended targets.

Here, we develop a strategy to confer controllable motility to bacteriophages by incorporating iron nanoparticles (FeNPs) into their head structures, enabling our magnetic guidance. Unlike genetic or chemical modifications that alter phage surface properties (Carmody et al., 2021), our method preserved the native infectivity of Fe-NP- tagged phages while introducing an external control mechanism for directed movement. The magnetically-responsive phages, herein termed as ‘magnetophage’, possessed the FeNPs in their head structures, where FeNPs were confirmed to be present by TEM and Cryo-EM. Our EDS confirmed that the NPs were iron in nature. The magnetophages could be first enriched and isolated using magnetic fields rapidly, eliminating the need for ultracentrifugation. Next, we further showed that magnetophages could be actively guided through complex environments, including microfluidic mazes, polymers and ex-vivo tissues, thereby bypassing diffusion constraints and enhancing targeted elimination of bacteria and their biofilms.

For *in vivo* application, we employed magnetophages to treat a Medaka fish tail wound infection model (Liu et al., 2024), as fish models are frequently used to study viral dissemination across host organs and tissues (Ayala-Nunez et al., 2019; Gaudin and Goetz, 2021; Zou and Nie, 2017). We could rapidly direct magnetophages from the injection site at the peritoneum to a distal infection site in the tail. This effectively eliminates the biofilm infection at the tail while avoiding off-target accumulation in other organs in the animals. Hence, our findings establish a new paradigm in virus-based therapies by introducing a controllable mechanism for viral transport, unlocking new possibilities for precision-targeted applications in medicine and biotechnology.

## Results

### Synthesis and Characterization of Magnetophages

We developed magnetophages by incorporating FeNPs into their head structures (**Fig. 1A**). Host *P. aeruginosa* PAO1 were first treated with FeNPs, so that they could incorporate FeNPs into their cytoplasm. Once the *P. aeruginosa* bacteriophage-2 (ATCC 14203-B1) (McVay et al., 2007) was added to infect the FeNP-tagged host bacteria, FeNPs in the bacterial cytoplasm were passively introduced into the phages’ head during phage synthesis, leading to the inclusion of FeNPs into the phages’ head. It is important to note that the FeNPs must be smaller than the phage’ heads, to ensure that the nanoparticles could be incorporated into the phages. Transmission electron microscopy (TEM) confirmed the successful integration of NPs into the phage capsids, without affecting the overall structure of the phage (**Fig. 1B**). If FeNPs were added directly to the native phages without the presence of the host bacteria, FeNPs were not incorporated into the phages (**Fig. 1B**). Our TEM-coupled EDS also confirmed that the NPs’ composition was primarily iron in nature, whereas the detected copper (Cu) mainly belonged to the metal grid (**Fig. 1C**). Despite possessing FeNPs in the head, the magnetophages maintained similar viabilities as their native phages, while FeNPs only did not impose any killing effect on the bacteria (**Fig. 1D**). We could enrich and isolate the magnetophages from lysed bacteria cultures into a visible pellet using a neodymium magnet (**Fig. 1E and Supplementary Video 1**), thus eliminating the need for ultraspeed centrifugation which is frequently used for hours to enrich phages (Dika et al., 2013). Moreover, our approach could be used for different phage species that target different bacterial species (*Stenotrophomonas maltophilia and Pseudomonas aeruginosa)*, where they could incorporate FeNPs and selectively kill their host bacterial species (**Supplementary** Fig. 1).

**Figure 1.**
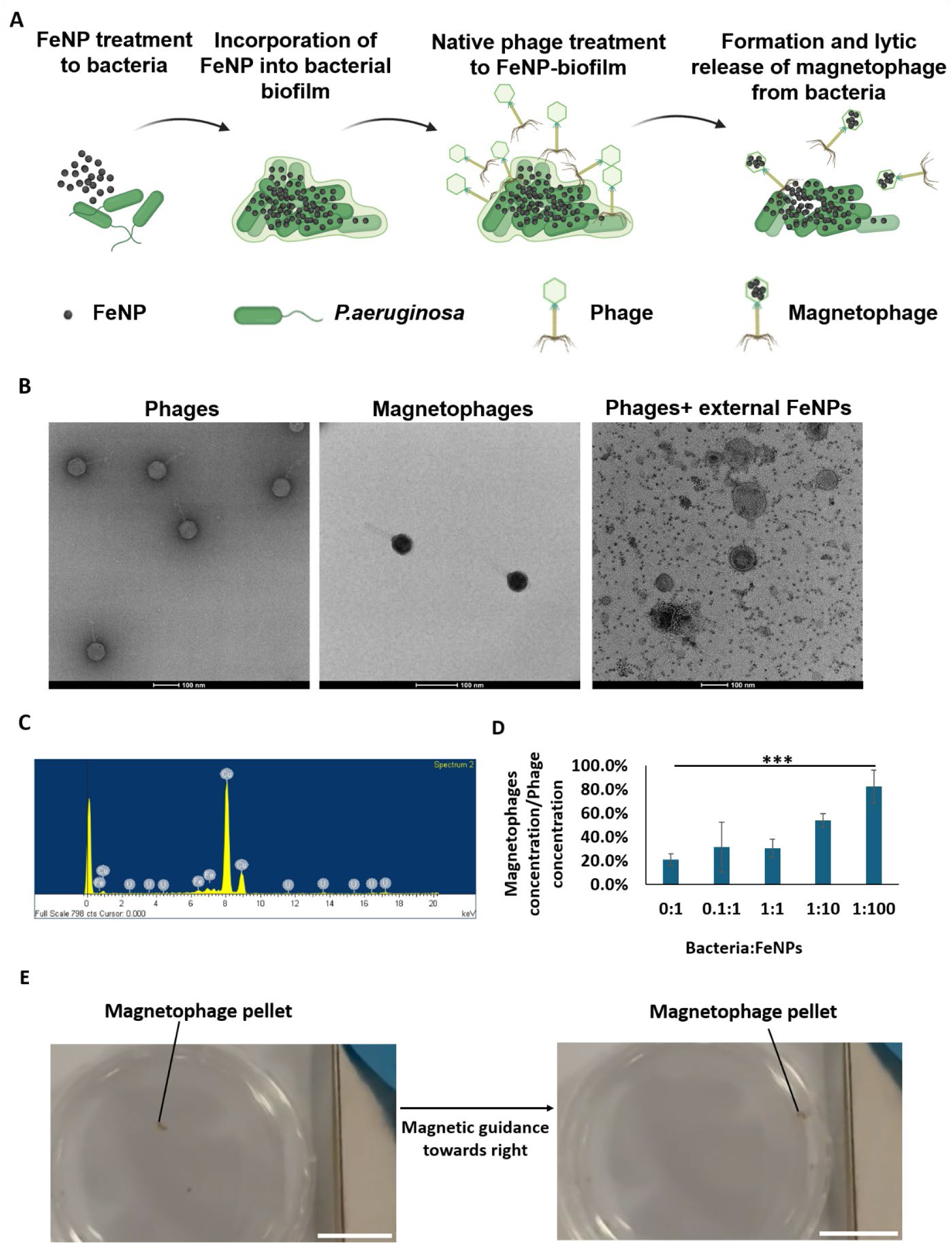
Synthesis and Characterization of Magnetophages. (A) Schematic diagram of incorporating iron nanoparticles into phages. (B) Representative TEM images of native phages and magnetophages. Scale bar: 100 nm. (C) EDS spectrum of magnetophages revealed presence of Fe. (D) Similar killing efficacy of magnetophages to native phages. (E) Representative photo of visible magnetophage pellet after isolating from bacteria. Scale bar: 1 cm. The means and s.d. from triplicate experiments from 3 independent trials were shown. ***p < 0.001.

### Cryo-EM confirmed the presence of FeNPs in the phage’s head structure

For high-resolution observation of FeNPs in the phage’s head structure, we employed cryo-EM to show that presence of individual FeNPs together with phage DNA (**Fig. 2A**). We observed a direct correlation between initial FeNP concentration and the number of FeNPs incorporated per phage, whereby adding lesser initial FeNP would result in fewer FeNPs incorporated into the phage (**Fig. 2B**). Hence, 80 μg/ml is the optimal FeNP concentration for introduction into phages (**Fig. 2C**). Interestingly, when the tail was lost from the phage, the FeNPs retained within the phage head (**Fig. 2D**). Since FeNPs cannot be transferred to the intented host, the newly synthesized phages would not possess any FeNPs and therefore their magnetic attribute after reaching their intended host and initiate the first infection (**Supplementary** Fig. 2). It was only when the phage head was damaged after phage lysis, then the FeNPs were released into the milieu (**Fig. 2E**). Hence, our cryo-EM images confirmed that the FeNPs could be incorporated into the phage’s head structure.

**Figure 2.**
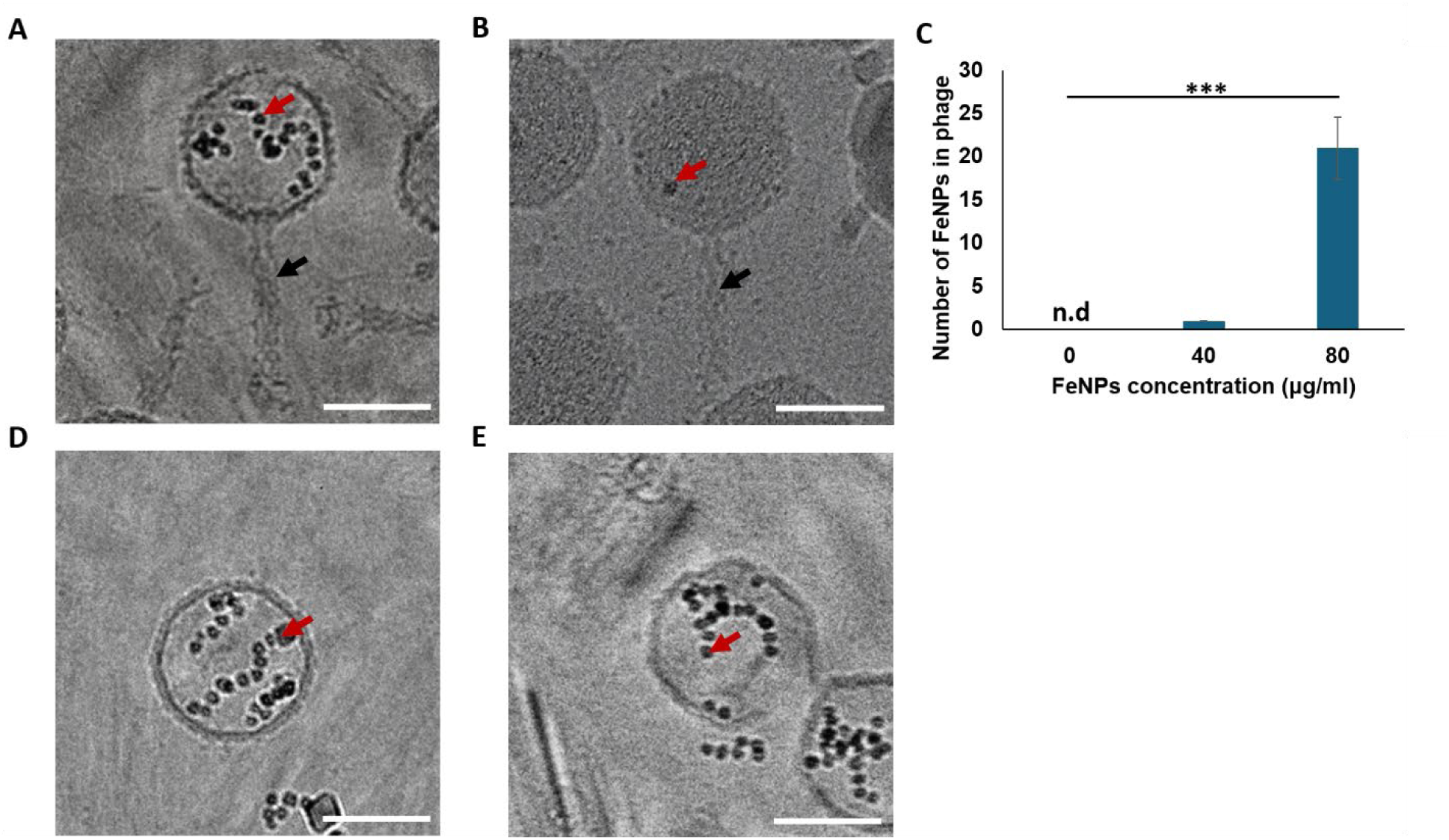
Cry-EM imaging of magnetophages. (A) Representative cryo-EM image of magnetophage with initial 80 µg/ml concentration of FeNPs added to biofilm. (B) Representative cryo-EM image of magnetophage with lower concentration of FeNPs added to biofilm. (C) Concentration of iron nanoparticles used for incorporation is correlated with number of iron nanoparticles in phages. (D) Representative cryo-EM image of magnetophage after loss of tail. (E) Representative cryo-EM image of a lysed magnetophage. Red arrow: FeNP. Black arrow: phage tail. Scale bar: 50 nm. The means and s.d. from triplicate experiments from 3 independent trials were shown. ***p < 0.001. n.d: not detected.

### Magnetic Control of Magnetophage Motility

We next evaluated whether magnetophages could be guided across distances using an external magnetic field. A microfluidic maze device with primary and secondary chambers was designed to test directional movement of magnetophages (**Fig. 3A-B**).

**Figure 3.**
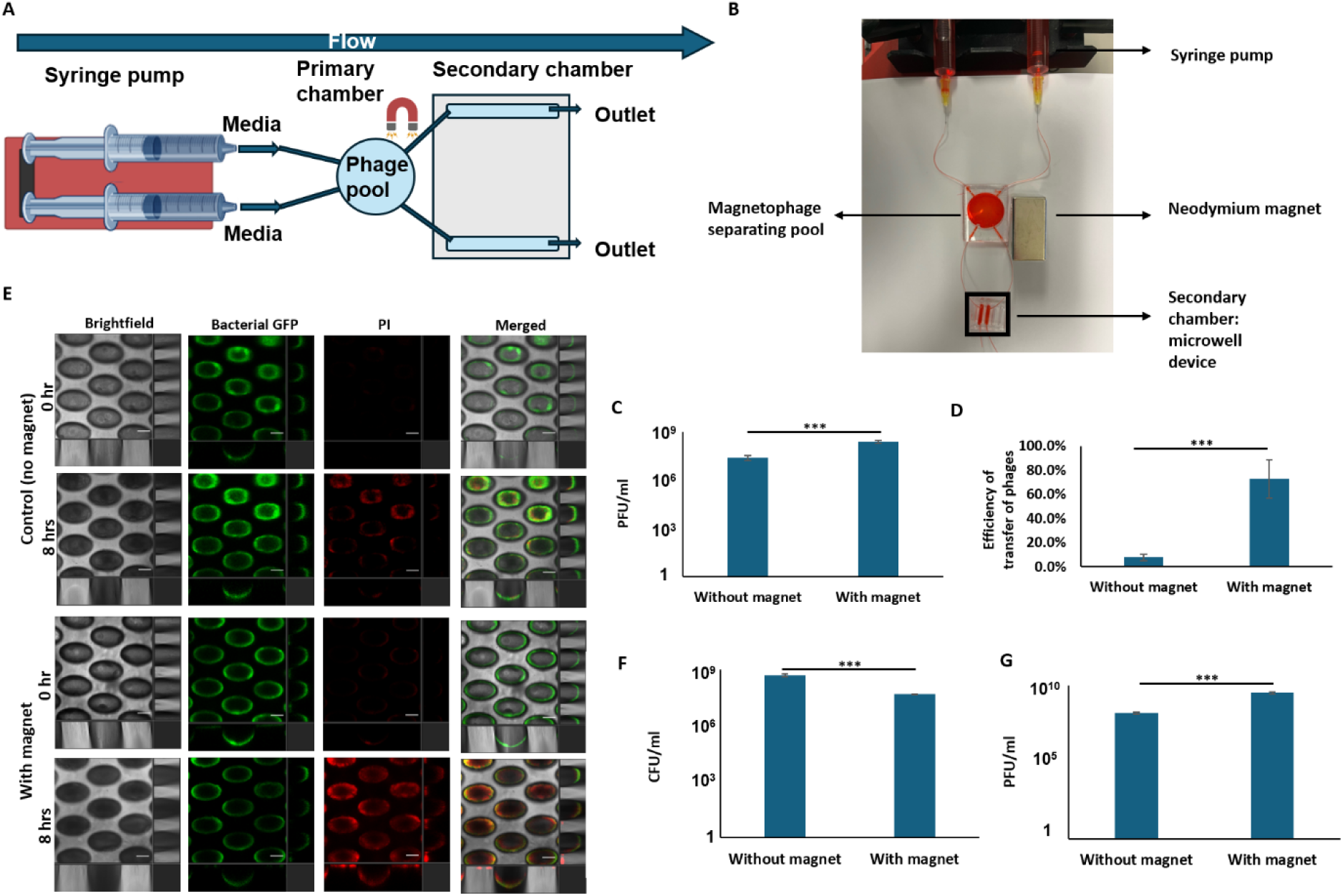
Magnetic Control of Magnetophage Motility. (A) Schematic diagram of microfluidic maze device with primary and secondary chambers. (B) Representative image of microfluidic device. (C) PFU of native and magnetophages in secondary chamber directly after magnetic manipulation. (D) Efficiency of transfer of magnetophages from primary to secondary chamber. (E) Representative CLSM images of biofilms in both control and magnetically controlled secondary chambers. Scale bar: 100 µm. (F) CFU of bacteria in both secondary chambers after 8 hrs incubation with phage treatment. (G) PFU of native and magnetophages in both secondary chambers after incubation. The means and s.d. from triplicate experiments from 3 independent trials were shown. ***p < 0.001.

Our PFU assay demonstrated that magnetophages accumulated in the secondary chamber upon magnetic stimulation (**Fig. 3C**), where the efficiency of transfer could reach more than 75% in 2.5% agar (**Fig. 3D**). When bacterial lawn was cultivated in the secondary chambers, the magnetophages could be directed towards the target bacterial lawn and eliminate their host (**Fig. 3E-F**), while the PFU of the magnetophages increased correspondingly (**Fig. 3G**). The non-targeted bacterial lawn remained uninfected by the magnetophages, with no detectable phages identified (**Fig. 3C-F**). In contrast, normal phages without FeNP tagging diffused and killed bacteria in both chambers (**Fig. 3C-F**). Hence, we showed that magnetophages could be magnetically manipulated to target sites.

### Penetration of Magnetophages into matrices

We next determined if magnetophages could penetrate through and traverse complex matrices, including tissues. Diffusion of native phages through tissues are typically slower than antibiotics, due to their larger sizes and distinct surface properties (Hu et al., 2010). To determine whether magnetophages could traverse matrices, we established a matrix penetration model using different concentrations of agar (**Fig. 4A**).

**Figure 4.**
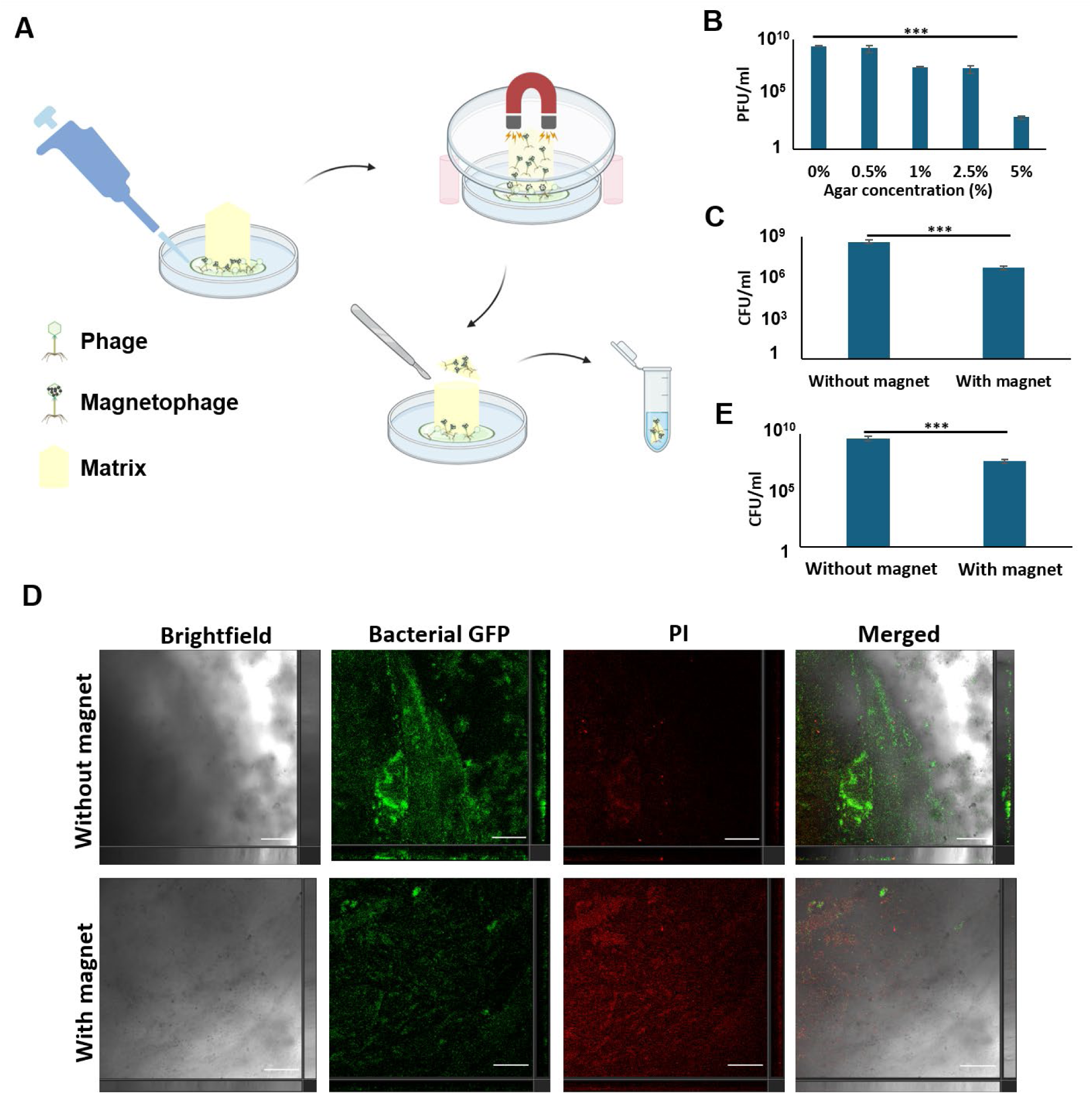
Penetration of Magnetophages into matrices. (A) Schematic diagram of phage penetration model using agar and *ex vivo* porcine skin wound model. (B) PFU of magnetophages and (C) bacterial CFU in the distal agar area at different agar concentrations. (D) Representative CLSM images of biofilms and (E) bacterial CFU in the wound infection site of the porcine skin model with and without magnetic treatment of magnetophages. Scale bar: 100 µm. The means and s.d. from triplicate experiments from 3 independent trials were shown. ***p < 0.001.

The magnetophages could migrate in the direction of the magnet through all agar concentrations, including up to 5% agar, where the phages could be detected (**Fig. 4B**) and bacteria at the distal bottom of the matrix were eliminated (**Fig. 4C**). In contrast, without magnetic direction, the magnetophages took significantly longer duration to diffuse through the softer agar, but could not pass through the hard agar (**Fig. 4B-C**).

Our results were corroborated with an adapted *ex vivo* porcine skin tissue (Gou et al., 2023) to show that the magnetophages could be magnetically transferred across the skin tissue to eliminate the bacteria located distally from the tissue (**Fig. 4D-E**). This demonstrated the effective penetration by magnetophages through various matrices.

### Anti-biofilm activity of Magnetophages against P. aeruginosa and its clinical isolates

Since the sticky matrix of mature and thick biofilms could also block the diffusion of phages to prevent elimination of its resident bacteria, or even promote further biofilm formation (Eriksen et al., 2018; Gödeke et al., 2011; Heilmann et al., 2012; Hu et al., 2012), we next assessed the antibiofilm efficacy of magnetophages against biofilms. Magnetophage treatment significantly reduced the colony-forming units (CFU) of wild-type PAO1 biofilms (**Fig. 5A**), while correspondingly increasing the phage numbers (**Fig. 5B**). Our confocal laser scanning microscopy (CLSM) showed that the biofilms were eliminated (**Fig. 5C**), where there was a decrease in GFP-labeled live bacteria (**Fig. 5D**) and an increase in dead bacteria stained by propidium iodide (PI) (**Fig. 5E**). Moreover, the magnetophages were effective against a carbapenem-resistant *P. aeruginosa* clinical isolate (PAE1004) (Ma et al., 2023a) and a cystic fibrosis patient-derived clinical isolate (CF173-2005) (Chua et al., 2016a) (**Fig. 5F-G**). This indicated that the magnetophages were effective at penetrating through the biofilms to kill the resident bacteria.

**Figure 5.**
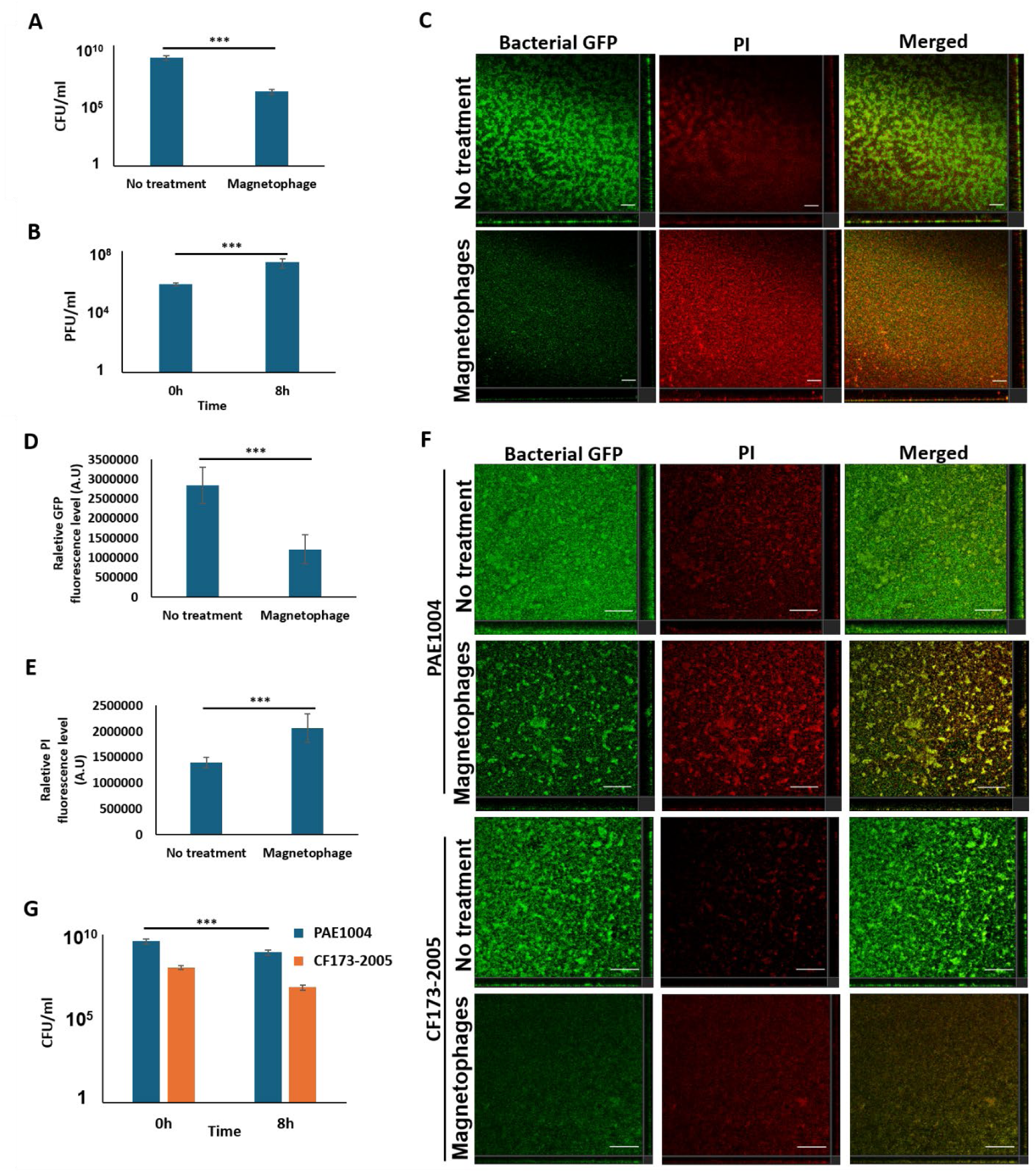
Anti-biofilm activity of Magnetophages against *P. aeruginosa* and its clinical isolates. (A) CFU of biofilms after native phage or magnetophage treatments. (B) PFU of phages after treatment of biofilms. (C) Representative CLSM images of biofilms. Scale bar: 50 µm. Relative (D) GFP and (E) PI fluorescence levels of biofilms. (F) CFU of *P. aeruginosa* clinical isolates’ biofilms after treatment with magnetophages (G) PFU of phages after treatment of clinical isolates’ biofilms. The means and s.d. from triplicate experiments from 3 independent trials were shown. ***p < 0.001.

### Targeted magnetophage treatment in vivo infection model

To assess if magnetophages could be employed as targeted therapy against localized infection with limited dissemination to other organs, we evaluated the *in vivo* therapeutic potential of magnetophages in a Medaka fish tail wound infection model (Liu et al., 2024) (**Fig. 6A**). Firstly, magnetophages used in our work at 10^6^ PFU/ml were not cytotoxic to human lung fibroblasts and Medaka fish animals (**Supplementary** Fig. 3). Magnetophages were successfully guided from the peritoneum to the *P. aeruginosa* biofilm-infected wound at the tail using the magnet (**Fig. 6B**). CLSM imaging confirmed the reduction of biofilm infection at the wound site (**Fig. 6C**), as indicated by decreased GFP signal (**Fig. 6D**). This was corroborated with lower bacterial counts (**Fig. 6E**), showing that bacterial burden was significantly reduced at the wound. Correspondingly, cytokine (TNF-α and IL-2) quantification of the fish tail displayed reduced inflammatory response (**Fig. 6F**). Moreover, to evaluate if the magnetophages were disseminated via the bloodstream to other body sites, we also collected liver samples and observed very low levels of magnetophages in the organ (**Fig. 6G**).

**Figure 6.**
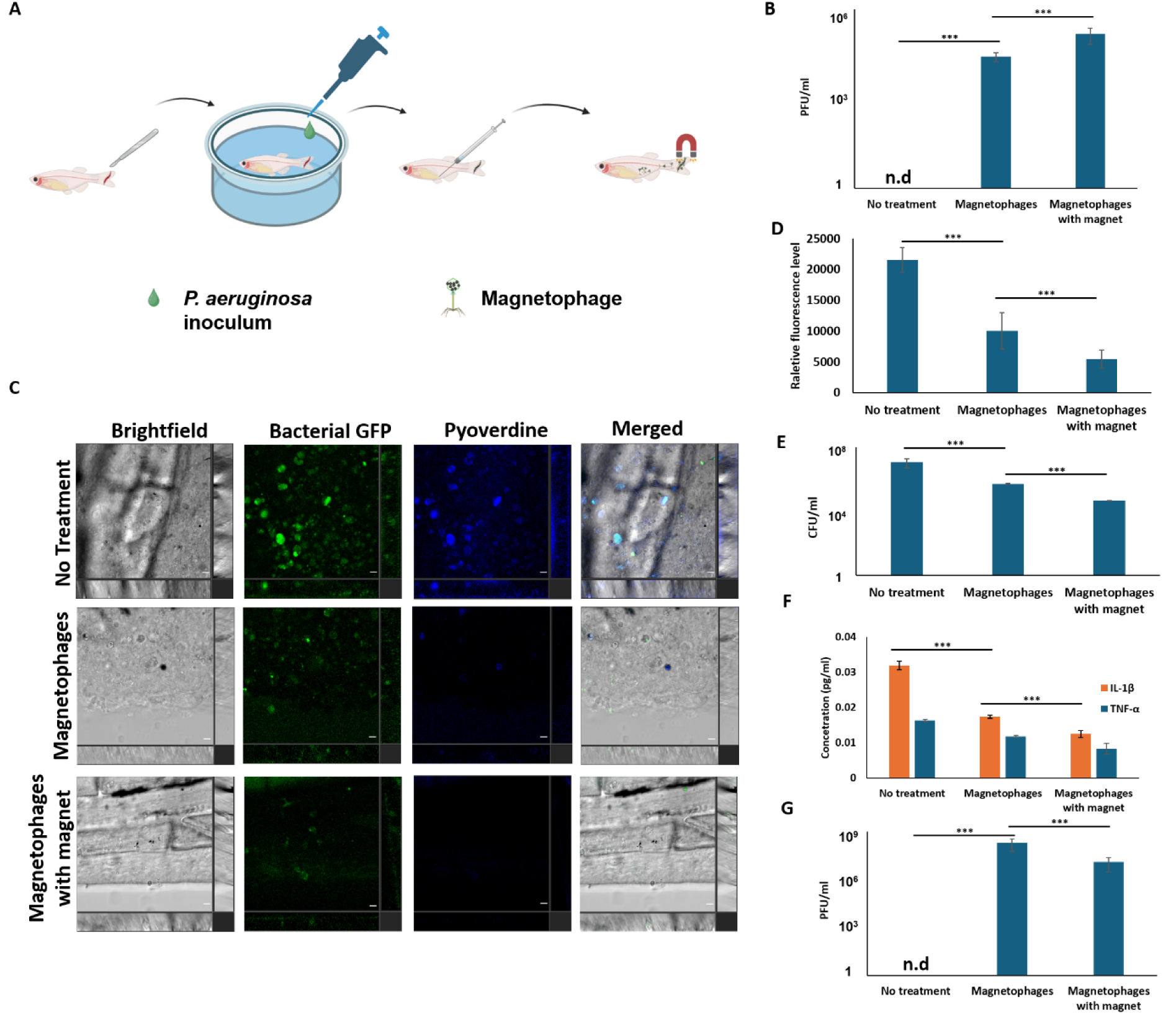
Targeted magnetophage treatment of fish tail infection model. (A) Schematic diagram of targeted antimicrobial treatment of fish tail infection using magnetophages. (B) PFU of magnetophages at tail wound. (C) Representative CLSM images of biofilms at tail wound. Scale bar: 10 µm. Relative (D) GFP and (E) PI fluorescence levels of biofilms at tail wound. (F) CFU of bacteria at tail wound after magnetophage treatment. (G) Relative cytokine (TNF-α and IL-2) concentration at the tail wound after magnetophage treatment. (H) PFU of magnetophages in the liver after magnetophage treatment. The means and s.d. from triplicate experiments from 3 independent trials were shown. ***p < 0.001. n.d: not detected.

In contrast, absence of magnetic manipulation of magnetophages would lead to lower numbers of phages at the site of infection, resulting in poorer clearance of biofilm infection and higher levels of inflammation (**Fig. 6B-G**). Furthermore, magnetophages were accumulated at higher levels in the liver, indicating widespread dissemination of phages to the other parts of the body (**Fig. 6G**).

## Discussion

Viruses, including bacteriophages, have traditionally been regarded as passive agents that rely on diffusion to reach their targets (Kaler et al., 2022). This fundamental limitation constrains the efficacy of virus-based therapeutic strategies, ranging from phage therapy to tumor virotherapy, as viral particles can accumulate nonspecifically throughout the body, leading to off-target toxic effects and suboptimal treatment outcomes (Lin et al., 2023). For phage therapy, phages face further problems in poor diffusion through high-density areas (Hu et al., 2010), including the tissues and high-density biofilms (Chegini et al., 2020), and may target resident bacteria in the host microbiome.

Here, we introduce a novel strategy to confer controlled motility to bacteriophages by passively incorporating FeNPs into their head structures, enabling external magnetic guidance and overcoming the diffusion limitations that often hinder phage therapy. Our findings demonstrate that magnetophages could be directed to navigate complex environments, penetrate biofilms, and selectively eliminate bacterial infections *in vivo*. Besides expanding the functional capabilities of viruses beyond their natural constraints, there are several advantages: [1] magnetophages could be targeted directly at the site of infection with limited off-target effects; [2] the loss of FeNPs from magnetophages after first infection of bacteria at the target site restores the native phages with minimal effects on humans; [3] our approach circumvents the need for chemical modifications that may compromise phage infectivity, preserving their natural ability to recognize and lyse specific bacterial hosts; and [4] our method reduces the tedious and expensive method to isolate and enrich phages for experimental or therapeutic purposes (Luong et al., 2020; Thurber et al., 2009).

The ability to guide viruses with external forces opens new frontiers in therapeutic applications. In the context of antimicrobial therapy, magnetophages offer a targeted approach to treating bacterial infections. Unlike antibiotics, which are systemically distributed (Nielsen et al., 2024) and contribute to resistance development (Naghavi et al.), magnetically guided phages can be concentrated at infection sites, enhancing efficacy while reducing collateral damage to the host microbiome. Beyond bacterial infections, the concept of controllable viral motility could be extended to other biomedical applications. For instance, oncolytic viruses could be engineered with magnetic properties to home in on tumor sites, increasing their therapeutic index and minimizing systemic exposure. Similarly, gene therapy viral vectors, including viral delivery systems for CRISPR-based genome editing, could be directed with precision to specific tissues, with reduced risks of unintended gene modifications. In synthetic biology, magnetically responsive viruses could serve as programmable tools for biomaterial assembly, biosensing, or targeted drug delivery. By establishing a framework for engineering motile viruses, our work paves the way for novel applications that leverage viral precision and controllability.

Our study also highlights key considerations for future development. The biocompatibility of FeNP-tagged viruses must be further explored across different host systems, including humans, to assess potential immune responses and long-term effects. Additionally, the size, composition, and surface chemistry of FeNPs could be fine-tuned to optimize viral mobility, retention, and clearance (de Montferrand et al., 2013). The combination of magnetic guidance with external stimuli, such as light or ultrasound, may further enhance the precision of viral delivery in complex biological environments. In conclusion, our work establishes a foundation for engineering motile viruses, transforming their role from passive carriers to actively guided therapeutic agents, with cross-disciplinary implications across microbiology, medicine, and biotechnology.

## Materials and Methods

### Bacteria and cultivation conditions

Bacterial strains used in this study are listed in **Table S1**. Bacterial strains were inoculated in 2 ml of LB medium containing appropriate antibiotics for plasmid maintenance at 37 °C with shaking at 200 rpm for 16 hrs. Luria-Bertani (LB) medium (Difco, Becton Dickinson and Company, USA) and ABTGC medium were used for cultivation and experiment of bacteria.

### Phage cultivation and extraction assay

The *P. aeruginosa* bacteriophage-2 (ATCC14203-B1, USA) was used in this study.To enrich bacteriophages, the target bacterial host was inoculated with the phage at a 10:1 (v/v) ratio in 10 mL of Luria-Bertani (LB) broth (Difco, Becton Dickinson and Company, USA). The mixture was incubated at 37 °C with shaking at 200 rpm for 12 hrs to facilitate phage propagation (Roach et al., 2017). Following incubation, the culture was centrifuged at 4000 rpm for 10 mins to remove bacterial cells and debris. The supernatant was then passed through a 0.22-μm pore-size syringe filter (BIOFIL, China) to obtain a cell-free phage-containing filtrate (Roach et al., 2017). The resulting filtrate was stored at 4 °C for subsequent plaque assays and further analysis.

### Magnetophage formation assay

To allow prey bacteria to first incorporate FeNPs in their cytoplasms, a water-dispersible 5-nm-diameter Fe₃O₄ NP solution (Xi’an Ruixi Biotechnology Co,. Ltd, China) was added to 1 mL of LB broth at a final concentration of 100 μg/mL, along with 1 μL of the bacterial culture. The mixture was incubated in a 24-well plate (SPL, Korea) and at 37 °C for 12 hrs to enable biofilm formation and incorporation of FeNPs into the bacterial cells. The presence of Fe₃O₄ enabled magnetic manipulation of the biofilm, where the neodymium magnet could attract the bacterial cells in the direction of the magnetic field.

To enable the phage predator to prey on bacterial biofilms and incorporate FeNPs, fresh LB containing phages at a final concentration of 1 × 10⁴ PFU/mL was added to the biofilms. The cultures were incubated at 37 °C for an additional 8 hrs without agitation.

Subsequently, the culture was gently resuspended and centrifuged at 4000 rpm for 10 mins, followed by filtering the supernatant through a 0.22-μm membrane filter to purify the magnetophage culture. The presence of Fe₃O₄ enabled magnetic manipulation of the magnetophages, where the neodymium magnet could attract the phage pellet in the direction of the magnetic field.

### Plaque-forming unit (PFU) assay

Phage titers were determined using the double-layer agar plaque assay (Waturangi, 2023). Serial tenfold dilutions of the phage-containing liquid were prepared in 0.9%(w/v) NaCl saline. The prey bacteria was added to 5 mL of molten soft agar (0.5% agar, maintained at ∼45 °C), gently mixed, and immediately poured onto LB agar plates (1.5% agar) to form a uniform bacterial lawn (Phumkhachorn and Rattanachaikunsopon, 2010). Next, each phage dilution was transferred onto the surface of the solidified top agar, followed by incubation of plates at 37 °C for 16 hrs. Clear zones (plaques) formed on the bacterial lawn were enumerated, and phage titers were calculated as plaque-forming units per milliliter (PFU/mL). The PFU/ml was tabulated by plaque number × dilution factor / volume.

### Colony-forming unit (CFU) assay

The bacteria samples were diluted serially in 0.9% (w/v) NaCl saline and then transferred to LB agar plates (Ma et al., 2023b). After incubation at 37 ◦C for 16 hrs, the colonies grown on the plates were enumerated. The CFU/ml was tabulated by colony number × dilution factor / volume.

### Transmission electron microscope- energy-dispersive spectroscopy (TEM-EDS)

Phage samples were first concentrated to 10^12^ PFU/ml, followed by negative staining of the phage samples using the 1% uranyl acetate. The phage images were captured by TEM at 100k magnification according to manufacturer instructions. The EDS report was generated by the TEM-EDS (JEM-2010, JEOL, Japan).

### Cryo-EM sample processing

The 3.5 µl phage sample was applied to glow discharged Quantifoil R2/2 300-square-mesh copper grids. After 15 s of incubation, the grid was blotted for 3 s at 4 ℃, 100% humidity to remove the excess buffer. The sample grid was then plunged frozen into liquid ethane using a Vitrobot Mark IV (Thermo Fisher Scientific, Germany). Cryo-EM images were captured using an FEI Glacios 200Kv cryo-TEM (Thermo Fisher Scientific) equipped Falcon 4i camera with a pixel size 0.94A at 150K times magnification. All selected grids holes with Z adjustment were selected for imaging, whereas holes with thick ice layers (the thick ice layers cause limited particles in the center of the hole) or obvious ice contamination were excluded.

### CLSM imaging

All microscopic images of bacteria and their biofilms were imaged by a Leica TCS SP8 MP Multiphoton/Confocal Microscope system to monitor brightfield, bacterial GFP and bacterial pyoverdine fluorescence in Z-stacks. For observation of dead cells, 5 µM propidium iodide (PI) was added to the culture. Images were processed with ImageJ. For quantification of fluorescence levels, the ImageJ was used and the formula used is corrected total cell fluorescence (CTCF) = integrated density - (area of selected cells X mean fluorescence of background readings).

### Fabrication of the microfluidics-based magnetic separation device

As previously described (Ma et al., 2023b), we designed a microfluidic continuous-flow model (length: 5 cm; width: 3 cm; height: 1 cm), containing an upstream phage pool (diameter: 2.5 cm) and 2 downstream straight channels (1 with neodymium magnet and 1 control without any magnet) for our study. Firstly, the microfluidic molds were silanized with trichloro (1H,1H,2H,2H-perfluorooctyl) silane (Sigma-Aldrich, Germany) for 16 hrs in a vacuum desiccator. The microfluidic device was fabricated using polydimethylsiloxane (PDMS), and prepared from a Sylgard 184 silicone elastomer kit (Dow Corning, USA). This involved thoroughly mixing the base resin and curing agent in a ratio of 10:1 by weight. Finally, the various layers were assembled and treated with plasma for 5 minutes at high RF level (700 mmtor), followed by heating in a 70 °C oven for ∼2 hrs.

### Microfluidics-based magnetic separation device for biofilm assay

As previously described (Ma et al., 2023b), bacterial biofilms was cultivated in 2 secondary chambers of the microwell-based microfluidic device for 12 hrs at 37 ℃, with continuous flow of ABTGC media at 4 ml/hr. The neodymium magnet was placed beside 1 secondary chamber to attract the magnetophages, whereas the other secondary chamber remained magnet-free. The magnetophages were injected into the primary chamber (phage pool), allowed to enter into the secondary chamber with magnetic guidance for killing of the biofilms.

### Agar penetration model assay

The filtered phage-containing supernatant was transferred into a Petri dish. An agar column (diameter: 3 cm; height: 2.5 cm) of various concentrations up to 5% was placed at the center of the dish, and a neodymium magnet was positioned above the top of the agar to attract magnetophages toward the top of the agar column for 30 mins. Subsequently, the top of the agar containing the concentrated magnetophages was excised and immersed in 0.9% NaCl solution. The agar was gently shaken in 0.9% NaCl saline solution for phage quantification using PFU assay.

### Ex-vivo porcine lung tissue penetration model

Porcine lungs were freshly cut into 5-cm slices with a thickness of 0.5 cm, and washed 3 times with sterile 1X PBS saline. The lung tissue was transferred to a new petri dish containing 10 ml Dulbecco’s modified Eagle’s medium (DMEM) (Life Technologies) supplemented with 10% fetal bovine serum (Gibco) for culture. A 20 μl 100X diluted overnight bacteria culture was added to the media for bacteria to colonize the bottom of the lung tissue for 1 hr, followed by replacement of fresh DMEM + 10% FBS. After the infection of lung tissue, 20 μl of magnetophage (final concentration of 10^6^ PFU/ml) was added to the top of the lung tissue for phage treatment for 6 hrs in 37 ℃, 99% humidity. The lung tissues were sectioned for CLSM imaging or homogenized for CFU assay.

### Human macrophage cytotoxicity assay

Human monocytes (ATCC U937) were cultivated with Dulbecco’s modified Eagle’s medium (DMEM) (Life Technologies) supplemented with 10% fetal bovine serum (Gibco) until 70% confluency for 96 h at 37 °C, 5% CO2, and 99% humidity. To differentiate the monocytes into macrophages, the U937 monocytes were plated in T25 cell culture flasks (SPL, South Korea) at a density of 7 × 10^5^ cells in each flask and treated with 100 ng/ml phorbol 12-myristate 13-acetate PMA (eBioscience, USA) for 72 hrs. The culture medium containing PMA was renewed every 24 hrs.

For experiment, 1 X 10^6^ cells/mL macrophages were first cultivated in each well of a 24- well microplate for 24 hrs at 37 °C, 5% CO_2_ and 99% humidity. The cells were then washed once with 1 X PBS and treated with 200 μL of DMEM containing phages for 24 hrs at 37 °C, 5% CO2, and 99% humidity. As previously described (Chua et al., 2016b), 5 µM PI was added to the cells to stain dead macrophages and subsequent observation by CLSM (20X objective). To evaluate total cell viability, the PI-stained dead cells and total cells were enumerated to calculate the macrophage survival rate. The formula used is total number of live cells/ total number of cells X 100%.

### Fish tail infection model

All experiments involving Medaka fish were conducted according to the ethical policies and procedures approved by the ethics committee of the City University of Hong Kong animal research ethics committee (reference number: AN-STA-00000071), and Hong Kong Polytechnic University Animal Subjects Ethics Sub-committee (case number: 23- 24/761-ABCT-R-GRF). All Medaka fish experiments were performed according to the guidance of the Department of Health in Hong Kong. Medaka fish were used to establish a tail wound infection model as previously described (Liu et al., 2022).

Prior to surgery, one-month-old fish larvae were anesthetized in 0.015% tricaine methanesulfonate (MS-222, Sigma-Aldrich, USA). A 100-μm skin wound was then established on the tail using a sterile 26G needle (Terumo, Japan) under a microscope. For infection with *P. aeruginosa* PAO1, the bacterial inoculum (OD600 = 1) was diluted 1:100 in sterile water and pre-incubated for 1 hr on the wound to facilitate bacterial colonization at room temperature. Subsequently, the animals were maintained in fresh water for 48 hrs at room temperature to allow development of the infection.

For phage treatment, 10⁶ PFU/ml of magnetophages were injected into the peritoneum of the fish. A neodymium magnet was then applied to the tail region to enable localised accumulation of magnetophage at the wound site for 20 mins. A non-magnet treated control was also used to prevent magnetosome motility at the wound site. For negative control, no magnetophages were added. After 4 hrs of phage treatment, the animals were euthanized by immersing the animals in an overdose of MS-222 (0.05 mM) for 30 mins. For direct observation of the wound infection, the fish tail was directly observed using CLSM with a 20x objective, where brightfield, GFP, and pyoverdine images were captured.

### Sample collection from tail and liver for CFU and PFU assays

The tail tissue was surgically removed from the euthanized fish and ground manually by pellet pestle in 100 μL of 0.9% NaCl in 1.5 mL microcentrifuge tubes until homogenization. The liver was surgically removed from the fish, placed in 100 μL of sterile 0.9% (w/v) NaCl saline and ground with a sterile grinding rod. As described in previous sections, the PFU and CFU assays were then performed to quantify phage and bacterial numbers respectively.

### Quantification of fish cytokines by ELISA

TNF-α and IL-1β were quantified from the homogenized fish tail tissue samples using the cytokine ELISA kit (Bosun, Jiangsu, China) according to manufacturer’s instructions, with measurement at OD 450 mm by the microplate reader (Tecan Infinite M1000 Pro, Switzerland) with normalization with protein concentration.

### Statistical Analysis

Experiments were performed in 3 independent trials, with triplicate samples. Averages and standard deviations were calculated using Microsoft Excel. The one-way ANOVA and Student t-test (paired) were calculated using Graphpad Prism, where appropriate. All data and figures are shown as the mean±s.d.

## Acknowledgements

This research is supported by Environment and Conservation Fund (84/2021), Health and Medical Research Fund (HMRF-23220372), Research Centre of Deep Space Explorations (BBFQ and BBCZ) and Pneumoconiosis Compensation Fund Board (PCFB- ZJN2).

---

**Supplementary Figure 1.**
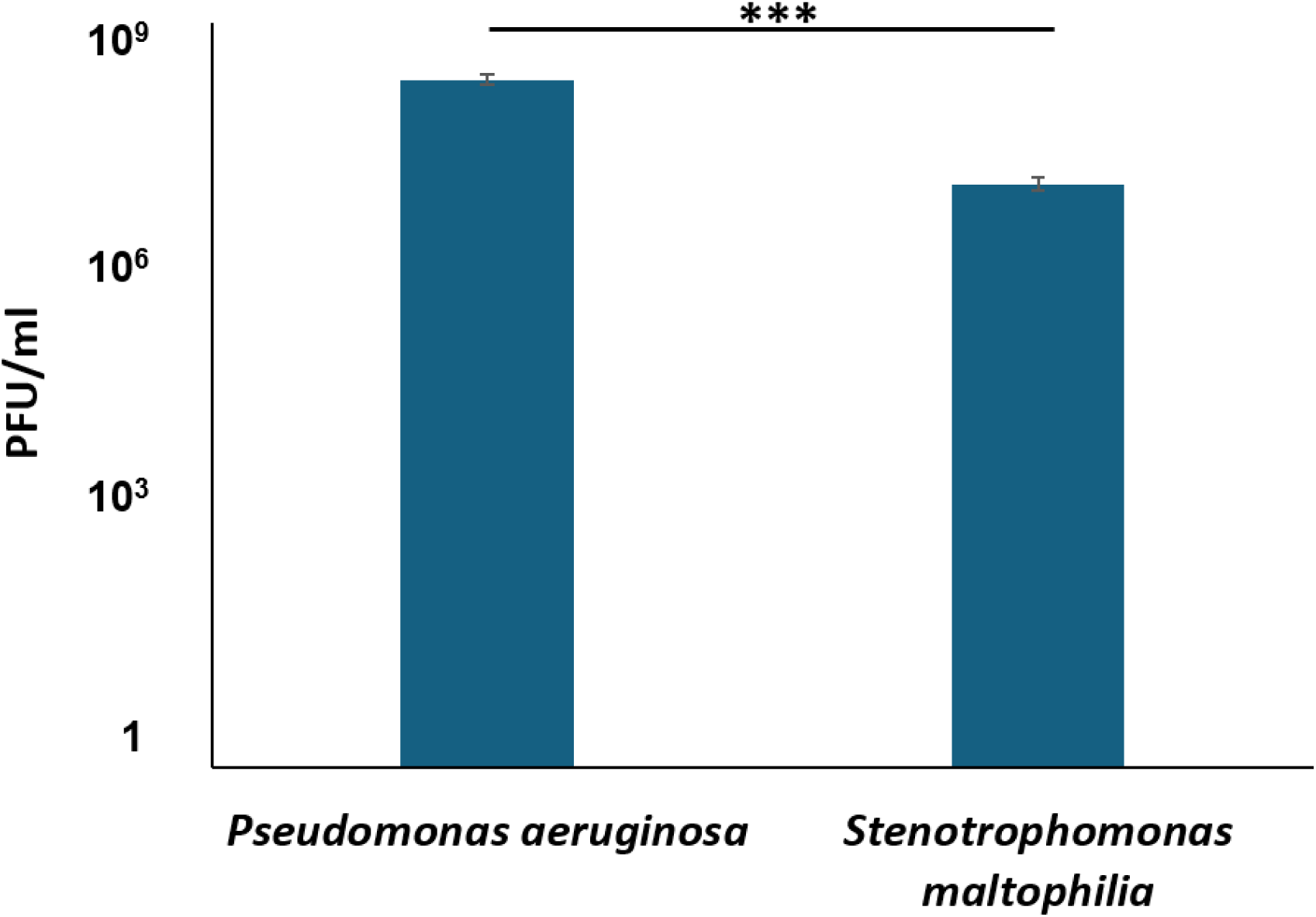
Different species of magnetophages can kill their target bacterial prey in the CFU assay. The means and s.d. from triplicate experiments from 3 independent trials were shown. ***p < 0.001.

**Supplementary Figure 2.**
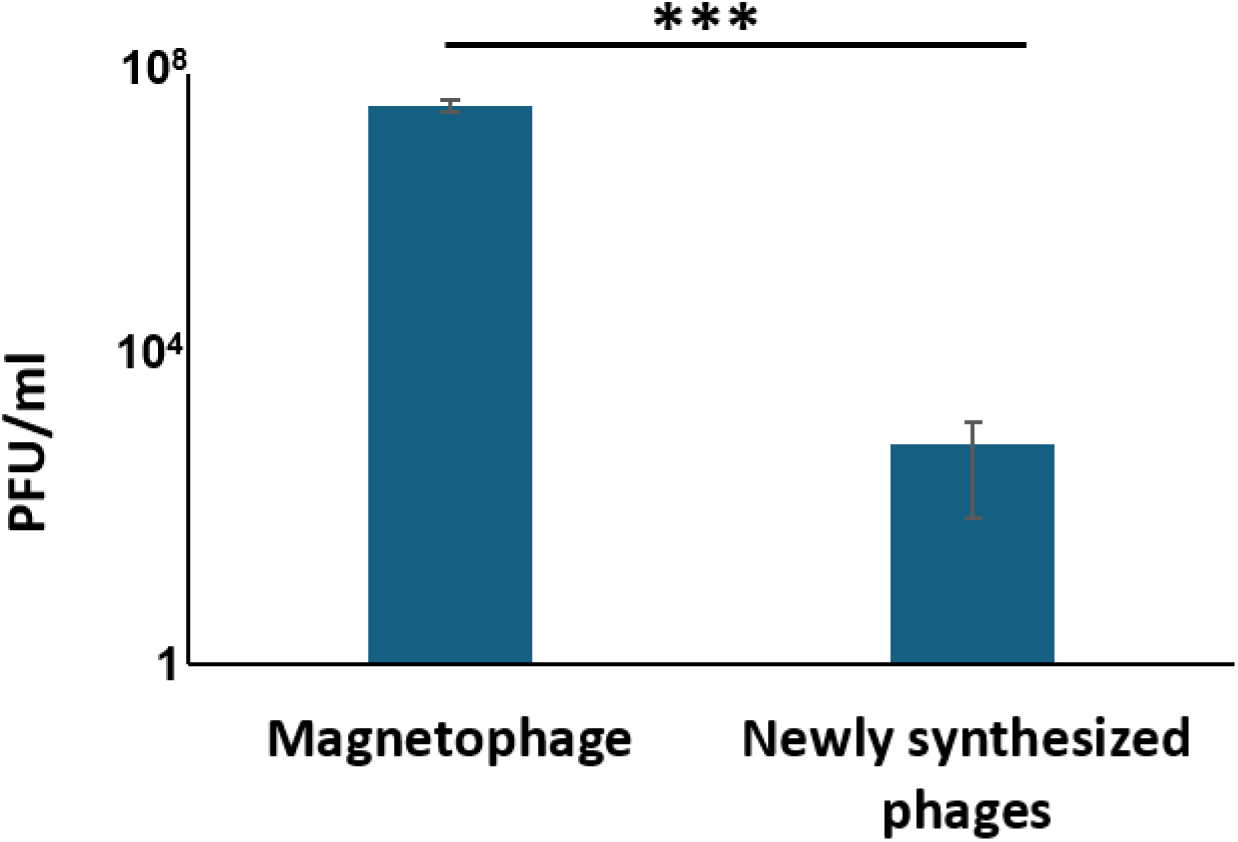
PFU of newly synthesized phages after subsequent infection of host. The means and s.d. from triplicate experiments from 3 independent trials were shown. ***p < 0.001.

**Supplementary Figure 3.**
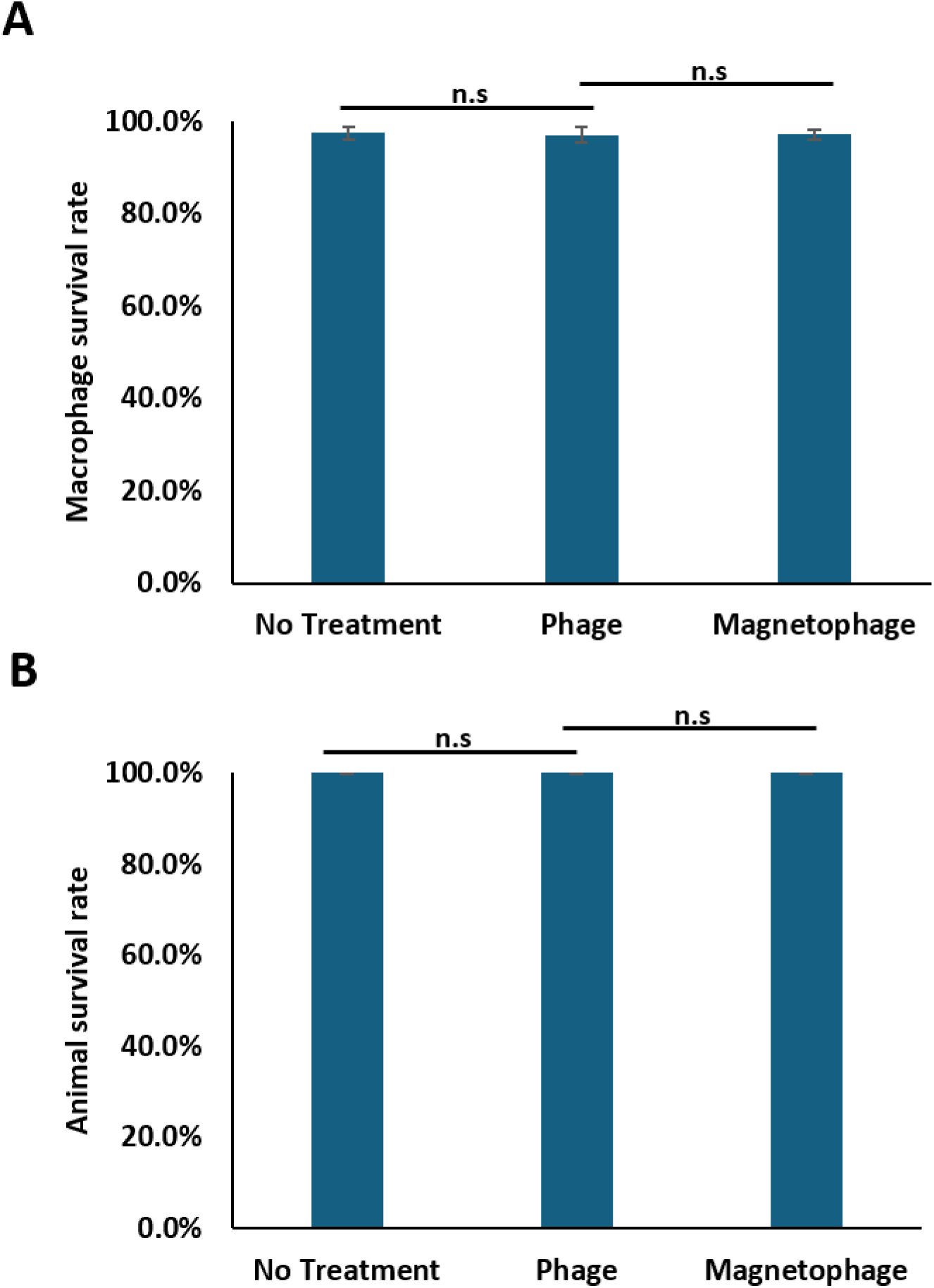
Magnetophages are non-cytotoxic to (a) mammalian cell cultures and (b) fish at our study-level concentrations. The means and s.d. from triplicate experiments from 3 independent trials were shown. N.s: not significant.

**Table S1.** Bacterial strains and plasmids used in this study.

| Strain/ plasmid | Description | Source/<br>Reference |
| --- | --- | --- |
| *P. aeruginosa* |  |  |
| PAO1 | Prototypic nonmucoid wild-type strain | (Holloway<br>and Morgan,<br>1986) |
| PAO1/*p<sub>lac</sub>-gfp* | Gm<sup>r</sup>, PAO1 containing the Tn7-transposon<br>(*p<sub>lac</sub>-gfp*), with constitutively expressed GFP | (Lambertsen<br>et al., 2004) |
| PAO1/*p<sub>cdrA</sub>-gfp* | Carb<sup>r</sup>, PAO1 carrying the c-di-GMP reporter<br>plasmid. | (Chua et al.,<br>2013) |
| PAE 1004 | Carbapenem-sensitive clinical isolate | (Ma et al.,<br>2023a) |
| CF173-2005 | Cystic fibrosis patient-derived clinical isolate | (Chua et al.,<br>2016a) |
| *S. maltophilia* |  |  |
| K279a | Bacteremia-associated clinical isolate | (Crossman<br>et al., 2008) |

**Supplementary Video 1.** Use of magnet to magnetically manipulate magnetophage pellet.

